# Confirmation of gamma irradiation–mediated inactivation of a Vero□adapted African swine fever virus Lisbon 60 strain for molecular assays

**DOI:** 10.64898/2026.05.20.726528

**Authors:** Sekhar Kambakam, Julia Thomas, Tod Stuber, Ping Wu, Suelee Robbe-Austerman, Rachel Palinski

## Abstract

African swine fever virus (ASFV), the etiologic agent of African Swine Fever (ASF), is a high-consequence pathogen requiring experiments to be conducted in containment in non-endemic countries, thereby restricting diagnostic development, the creation of reference standards, and proficiency testing (PT). Safe and reliable inactivation methods are essential to expand diagnostic capacity while preserving nucleic acid integrity for molecular assays in unaffected countries. This study employed gamma irradiation to achieve complete inactivation of ASFV without compromising downstream molecular detection, as gamma irradiation offers deep penetration and uniform dose delivery. ASFV-cell culture supernatants were subjected to gamma irradiation doses ranging from 2 to 50 kGy. Viral replication was evaluated using TCID□□ and serial passages, revealing a consistent dose□dependent reduction in infectivity across increasing irradiation dose levels and a complete loss of ASFV infectivity at 30 and 50 kGy. Molecular detection remained unaffected at all of the tested doses as confirmed by qPCR Ct values and sequence identity of the *p72* gene. Whole genome sequencing demonstrated >99% genome coverage and consistent read depth profiles across irradiated and non-irradiated samples, indicating preservation of genomic integrity at all tested doses. These findings demonstrate that gamma irradiation at 50 kGy fully inactivates ASFV-cell supernatants while maintaining nucleic acid quality suitable for molecular diagnostics. The resulting inactivated material meets quality assurance requirements for molecular reference standards and PT panels and can be safely distributed to laboratories outside high containment facilities, supporting broader diagnostic readiness and harmonization of ASFV testing.

## Introduction

African swine fever virus (ASFV) is a large, enveloped, double□stranded DNA virus belonging to the family *Asfarviridae* (Chapman et al., 2011; Alonso, 2018; Blome et al., 2020). ASFV continues to spread globally in domestic and wild swine populations, causing severe economic losses and posing major threats to animal health, food security, and international trade. African swine fever (ASF) is highly contagious and often fatal, with mortality rates reaching up to 100% in infected herds (Salguero, 2020; Blome et al., 2020). Currently, there are no licensed vaccines or treatments available (Burton et al., 2024), making rapid detection, expanded surveillance, and timely outbreak response essential for disease control (Burton et al., 2024; Kambakam et al., 2026).

Diagnostic testing for ASFV relies primarily on molecular assays such as qPCR (Pikalo et al., 2022a). However, ASFV is a high□consequence pathogen requiring BSL□3 or higher containment in unaffected countries (Pavone et al., 2024), limiting the number of laboratories and personnel capable of handling infectious virus. These biosafety constraints have also restricted the availability of standardized ASFV reference materials and proficiency□testing (PT) panels, hindering national and international diagnostic harmonization. Therefore, safe, effective, and rapid inactivation methods are needed to enable ASFV samples to be widely distributable as reference materials, for PT panels or diagnostic assay development, and other downstream applications in lower□containment laboratories.

Reliable and efficient inactivation methods are critical for high□risk pathogens such as ASFV. Inactivation methods, such as chemical (Brown, 1995; Blow et al., 2004; Haddock et al., 2016; Welch et al., 2024) and thermal (Woese, 1960; Delpuech et al., 2022) treatments, were shown to have adverse effects on the biological properties of pathogens, including antigenicity and nucleic acid integrity (Fan et al., 2015; Afrough et al., 2020; Pan et al., 2020; Elveborg et al., 2022). In contrast, ionizing radiation methods preserve structural and genomic features more effectively (Afrough et al., 2020; Elliott et al., 1982; David et al., 2017; Boudarkov et al., 2016; Feldmann et al., 2019; Elveborg et al., 2022; Leung et al., 2020). Although X□irradiation has been shown to inactivate multiple viruses while maintaining genome structure (Afrough et al., 2020), it has limited penetration and reduced dose uniformity in large□volume or high□density samples (Shahi et al., 2021). Gamma irradiation, typically generated from cobalt□60 (Co□60), provides deeper penetration and highly uniform dose delivery, making it well suited for inactivating dense or packaged biological materials (Shahi et al., 2021). Gamma irradiation has been widely used to inactivate a broad range of enveloped and non□enveloped viruses, while preserving structural and genomic integrity for molecular, serological, and biochemical assays (Elliott et al., 1982; David et al., 2017; Boudarkov et al., 2016; Feldmann et al., 2019; Elveborg et al., 2022; Leung et al., 2020; Pikalo et al., 2022b).

Although previous studies have demonstrated ASFV inactivation by gamma irradiation (McVicar et al., 1982; Boudarkov et al., 2016; Pikalo et al., 2022b), they have not fully characterized the suitability of inactivated material for downstream applications such as molecular diagnostic testing. Validating the gamma irradiation dose required to achieve ASFV inactivation while preserving genome integrity is essential for the production of standardized, biosafe ASFV reference materials and PT panels.

In this study, we aimed to determine the irradiation dose required for full inactivation of ASFV in infected cell□culture supernatant. We further sought to validate and confirm the absence of viral replication in irradiated ASFV through *in vitro* cell□culture assays. Additionally, we demonstrated that inactivated ASFV samples remain suitable for molecular testing without the loss of nucleic acid integrity, supporting their use in the preparation of ASFV reference materials and PT panels.

## Materials and Methods

### Virus and cell culture

The ASFV Lisbon 60 Vero cell-adapted strain (ASFV-L60V) was used in this study (Valdeira and Geraldes, 1985; Valdeira et al., 1998; Masujin et al., 2021), which was provided by the - U.S. Department of Agriculture (USDA) Foreign Animal Disease Diagnostic Laboratory (FADDL). The ASFV *in vitro* cell culture work was conducted at the biosafety level 3 facility at the USDA National Centers for Animal Health (Ames, IA, USA) in compliance with USDA and U.S. Department of Health and Human Services safety regulations regarding select agents.

ASFV was cultured in Vero cells using minimum essential media (MEM) supplemented with 5% fetal bovine serum, 4 mM L-glutamine (Gibco), 0.015 g/L of penicillin, 0.1 g/L of streptomycin, and 1x non-essential amino acids (Gibco) at 37 °C, 5% CO□ for 5 to 7 days. The virus-cell supernatant was clarified by centrifugation at 1500 g for 10 min at 4°C. The 50% tissue culture infectious dose (TCID_50_) was used for viral titering and calculated by the Reed-Muench method (Abolaban and Djouider, 2021). The clarified virus was concentrated for gamma irradiation using an Amicon 50 kDa cutoff centrifugal filter (Millipore) at 1500 g centrifugation at 4°C.

### Gamma irradiation facility

The gamma irradiator facility was established in compliance with Nuclear Regulatory Commission (NRC) and USDA regulations at a designated, secured, and access-controlled non-containment location. The irradiator used was a GammaCell 220F (Foss Therapy Services, CA, USA) equipped with a Co-60 radiation source.

### ASFV cell□culture supernatant for irradiation

ASFV infected cell culture supernatant was diluted to a titer of 10^6^ TCID_50_/mL, and 1 mL aliquots created in 2 mL cryogenic vials (ThermoFisher, MA, USA) or 20 mL aliquots were created in 35 mL Oak Ridge polypropylene tubes (ThermoFisher, MA, USA). We prepared sample packets consisting of ten 2□mL tubes or five 35□mL tubes, which were irradiated separately. Sample packets were placed in a canister with dry ice and irradiated at 2, 4, 7, 10, 30, and 50 kGy doses. Following irradiation, alanine dosimeters were removed and analyzed by GEX Corporation (https://www.gexcorp.com/) to determine the absorbed dose.

### Gamma irradiation ASFV sample validation in cell culture

The presence of competent ASFV in irradiated and non□irradiated ASFV was assessed using TCID_50_ and three blind serial passages in Vero cells. Virus samples were subjected to 1:10 serial dilutions and inoculated onto 70–90% confluent Vero cells in 96□well plates. Cultures were monitored for cytopathic effect (CPE) for 5–7 days, and TCID_50_ was calculated. For serial passages, virus samples from irradiated tubes and the non□irradiated controls were inoculated separately onto 70–90% confluent Vero cells in T□25 flasks at a 1:10 inoculation ratio. From the 20□mL irradiated virus in a 35□mL tube, we analyzed 12% of the total volume. Passages were collected and clarified by centrifugation at 1,500g for 10 minutes. The 1:10 diluted virus cell supernatant was added to fresh cells for each subsequent passage.

### Determination of the D_10_ value and sterilization dose

The sterilization dose (SD) required to achieve the internationally accepted sterility assurance level (SAL) of 10□□ can be determined using the D_10_ value (Gazsó and Gyulai, 2004; Fairand et al., 2010; Hewitt and Leelawardana, 2014). The D_10_ value means the radiation dose required to reduce 90% of viral infectivity (i.e., 1-log_10_). It was calculated as the negative reciprocal of the slope of the regression line generated from the log kill curve, which plotted viral titer against absorbed irradiation dose.

The SD was calculated using the initial viral titer (N), the D_10_ value, and the SAL of 10^-6^, according to the following equation (Gazsó and Gyulai, 2004; Fairand et al., 2010; Hewitt and Leelawardana, 2014):

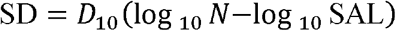

This calculation provides the irradiation dose required to ensure inactivation of ASFV to the accepted sterility standard.

### Nucleic acid extraction and qPCR

Nucleic acid (NA) extraction for qPCR and next□generation sequencing (NGS) was performed using the MagMax™ CORE Nucleic Acid Purification Kit (ThermoFisher, MA, USA) following the previously published protocol (Kambakam et al., 2026). Quantitative PCR (qPCR) assays were conducted according to the National Animal Health Laboratory Network (NAHLN) standard operating procedures, as described in Kambakam et al. (2026) and Zsak et al. (2005).

### Next generation sequencing (NGS)

The extracted NA concentration was determined using the Qubit dsDNA High Sensitivity Range Assay kit (Invitrogen, Carlsbad, CA, USA) on a Qubit Flex. Samples were prepared for sequencing on the Illumina Nextseq 2000 platform using the Illumina DNA library prep kit and sequenced using 2×150 P1 paired-end SBS (sequencing by synthesis) chemistry flow cell per manufacturer’s instructions (Illumina, San Diego, CA, USA).

### Genome coverage analysis

Genome coverage analysis was performed using paired-end Illumina sequencing reads from non-irradiated and irradiated (30 and 50 kGy) samples aligned against a reference genome (NC_044944.1). The analysis pipeline utilized Burrows-Wheeler Aligner (BWA-MEM) for read alignment, followed by coverage depth calculation using SAMtools (Li et al., 2009). Coverage statistics were computed across the genome, including percent coverage and average read depth in covered regions. Comprehensive visualizations were generated showing coverage patterns across the largest contigs, with comparative coverage plots displaying the sequencing depth distribution for all three samples (0, 30, and 50 kGy).

The *p72* gene target sequences from qPCR were aligned between non-irradiated and irradiated samples using Geneious Prime 2026.0 (https://www.geneious.com).

## Statistical analysis

The standard error of the mean (SEM) was calculated from three independent replicates. For statistical significance, unpaired 2-tailed Student t-tests were performed (Prism 10 software; GraphPad).

## Results

### Determination of the irradiation dose required to inactivate ASFV

ASFV□infected cell culture supernatants with an initial titer of 10^6^ TCID_50_/mL were exposed to gamma□irradiation doses of 2, 4, 7, 30, and 50 kGy. Absorbed doses were measured using alanine dosimeters, whose values (difference ±0–2) were consistent with the calibrated doses except at 50 kGy (Table 1), likely due to prolonged exposure to dry ice, as alanine dosimeters are known to be temperature□sensitive (Desrosiers et al., 2004).

**Table 1.**
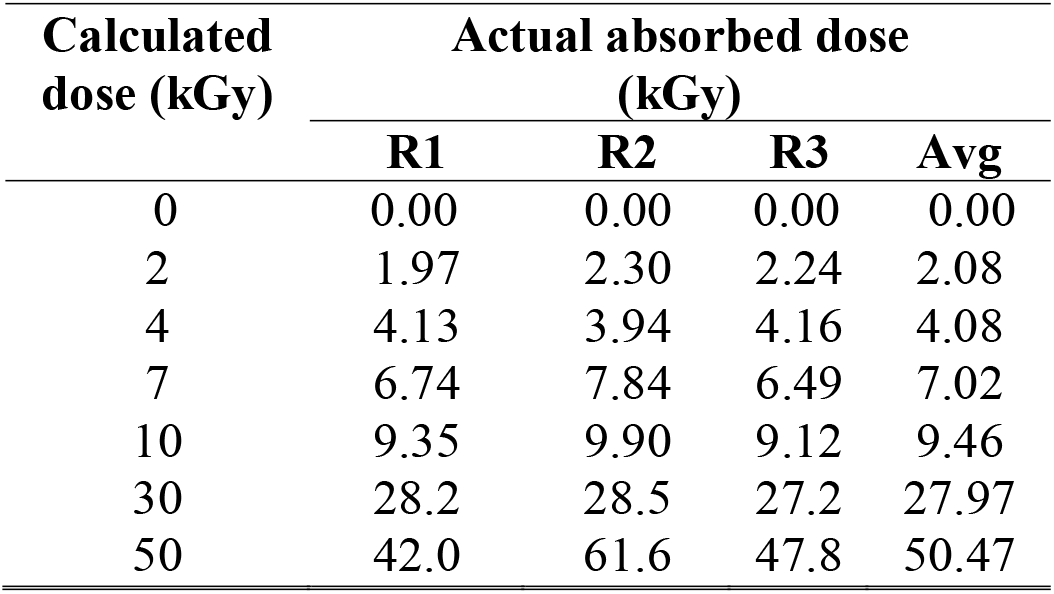
The actual absorbed dose received by the sample was calculated using irradiated alanine dosimeters. R1, R2 and R3 indicate three replicates.

Samples irradiated at 30 and 50 kGy exhibited complete loss of viral infectivity and were therefore not used to generate the kill curve. A linear relationship was observed between absorbed dose and reduction in viral titer (R^2^ = 0.9610) (Fig. 1). Based on this regression, the D_10_ value for ASFV was calculated to be 2.3 kGy. The calculated SD dose for ASFV was 29.1 kGy, indicating that exposure to approximately 29 kGy meets the internationally accepted sterility assurance level (SAL).

**Fig. 1.**
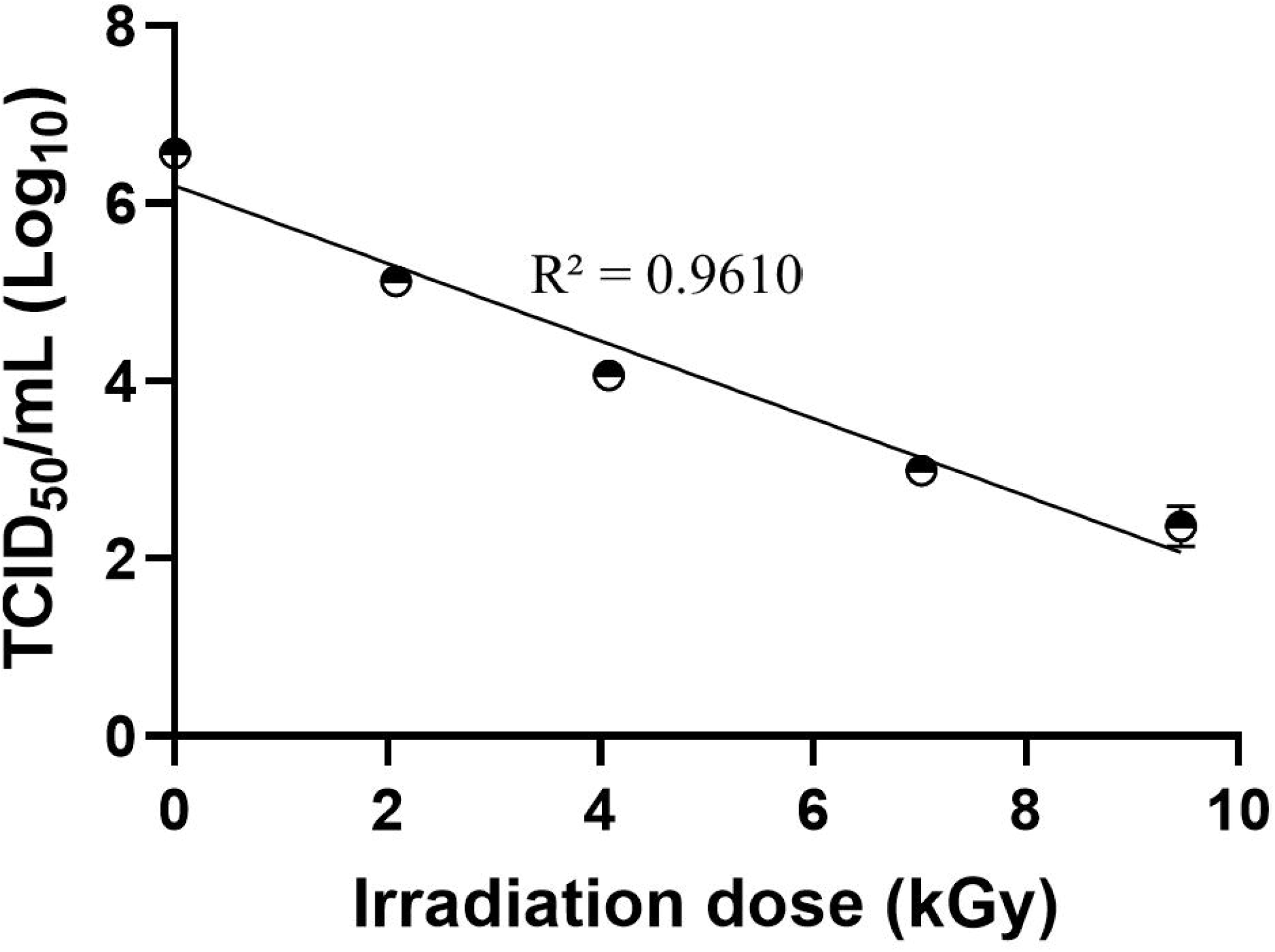
The log-kill curve was used to calculate the D_10_ value that is required to determine the gamma radiation dose to inactivate the ASFV. The data are x□ □±□ SEM from the 3 replicates.

### Effect of 30 and 50 kGy irradiation on ASFV infectivity *in vitro*

To ensure the virus was completely inactivated with 30 or 50 kGy, irradiated and non□irradiated samples were assessed by TCID_50_ assay using three blind serial passages on Vero cells. No CPE or viral replication was detected in samples irradiated at either 30 or 50 kGy. Conversely, as expected, virus replication was observed in non-irradiated samples (Fig. 2, Table 2). These results confirmed that ASFV was inactivated at both irradiation doses, consistent with the determined inactivation dose (~29 kGy) based on a linear log□kill curve.

**Fig. 2.**
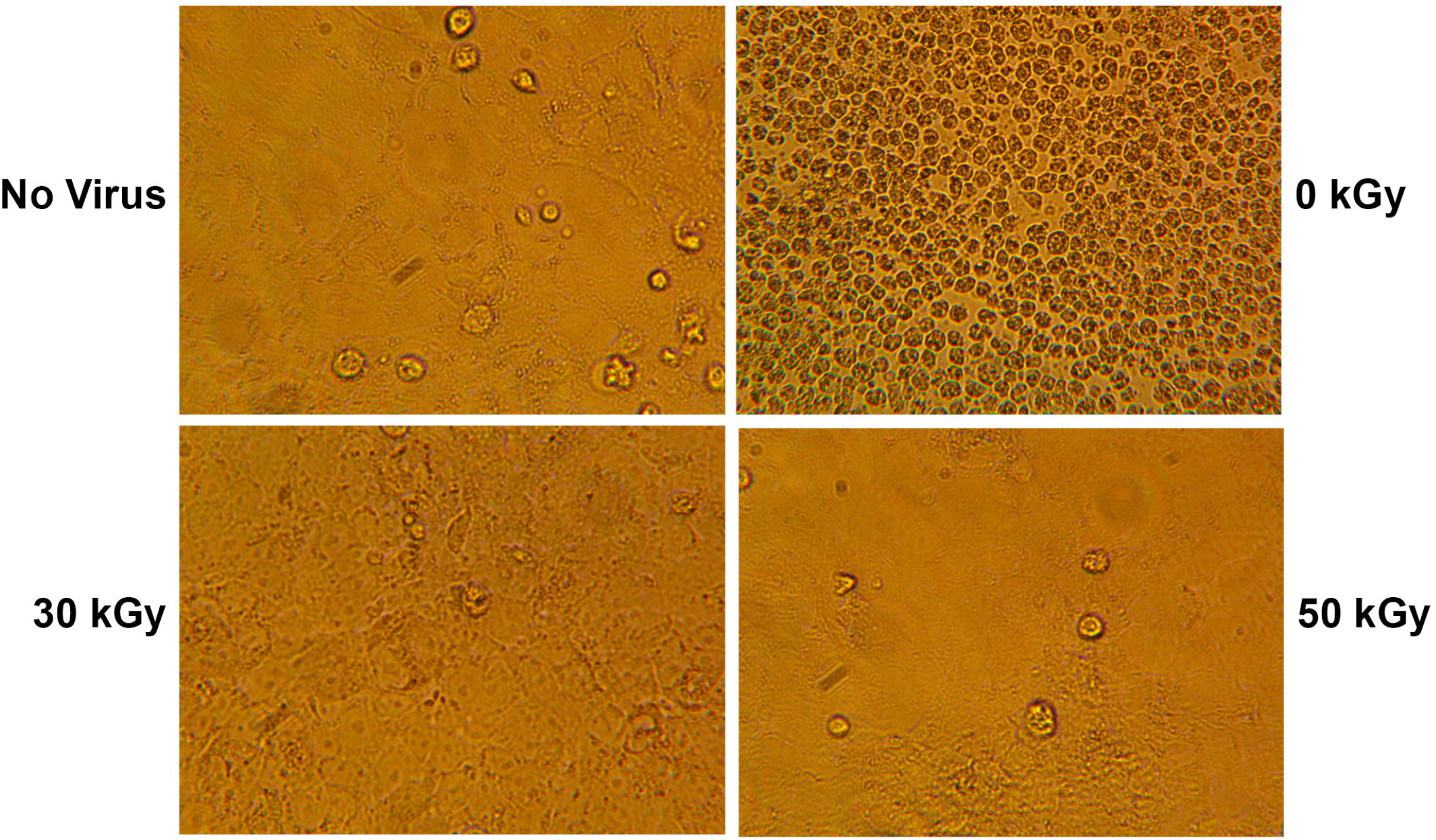
Visualization of CPE after serial passage of ASFV. The images represent one replicate from three independent replicates. Images were taken at 25x magnification. The data are x□ □±□ SEM from the 3 replicates.

**Table 2.** Observation of cell infectivity after adding gamma-irradiated ASFV.

| Dose (kGy) | Virus growth (P1/P2/P3) |
| --- | --- |
| 0 | +/+/+ |
| 2 | +/+/+ |
| 4 | +/+/+ |
| 7 | +/+/+ |
| 10 | +/+/+ |
| 30 | -/-/- |
| 50 | -/-/- |

‘+’ represents CPE; ‘-’ represents No CPE; P represents Passage

### Effect of high irradiation doses on the molecular detection of ASFV

To determine whether high-dose irradiation (30 and 50 kGy) affects molecular testing, qPCR Ct values for irradiated samples were comparable to those of non-irradiated samples, indicating that neither dose compromised qPCR detection of ASFV (Fig. 3A). Further, this result was confirmed by sequencing the qPCR target region between irradiated and non-irradiated samples resulting in 100% identity across all sequences (Fig. 3B). The third blind serial passage was tested by qPCR resulting in Ct values near the negative threshold, consistent with the absence of viral replication in cell culture (Fig. 4). In contrast, non-irradiated samples showed lower Ct values (Fig 4).

**Fig. 3.**
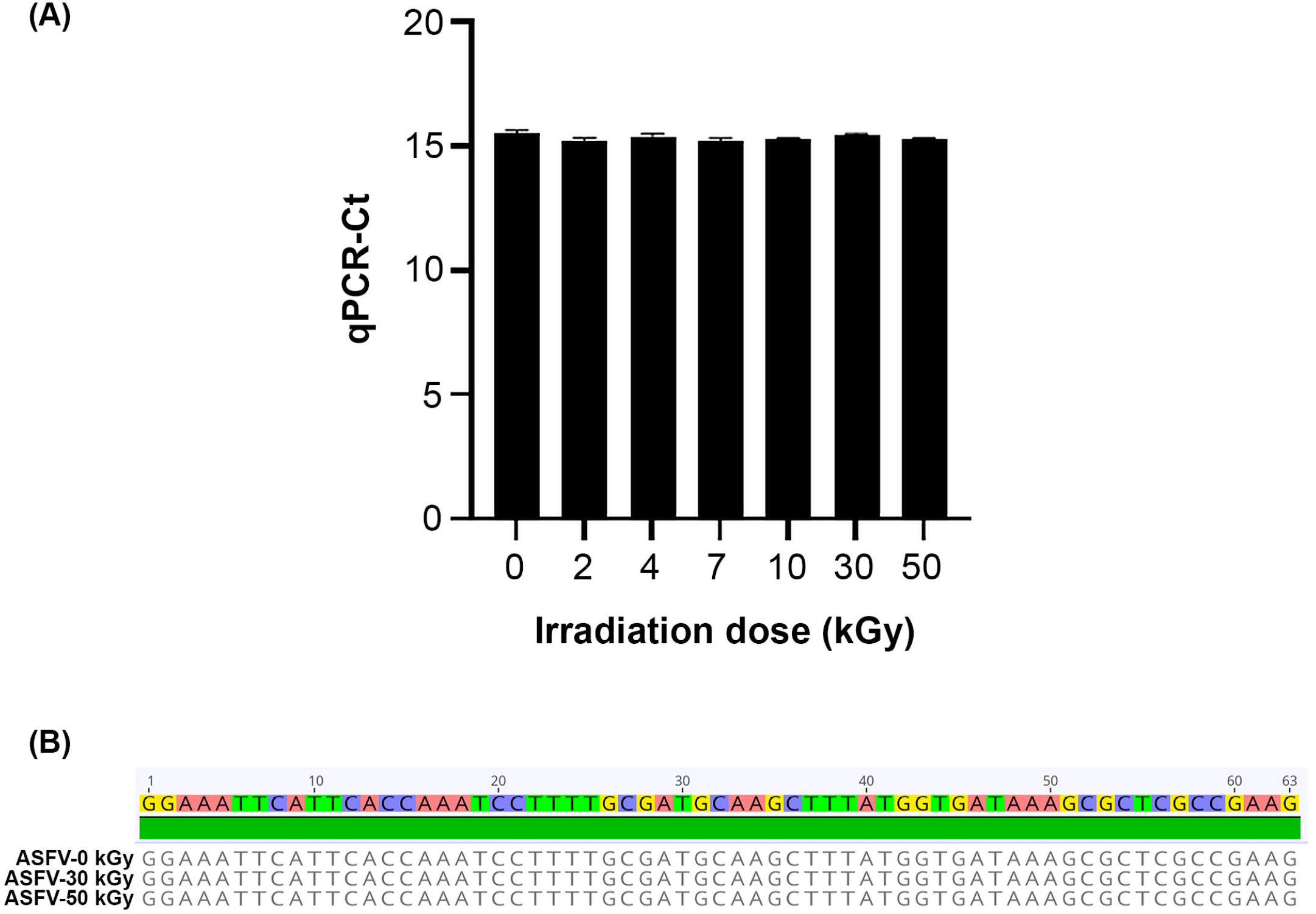
Assessment of nucleic acid integrity in gamma□irradiated samples for ASFV detection by qPCR. **A**. The qPCR Ct values of irradiated samples were the same as non-irradiated samples. **B**. The qPCR target sequence of *p72* gene aligned between non-irradiated (0 kGy) and irradiated samples (30 kGy and 50 kGy).

**Fig. 4.**
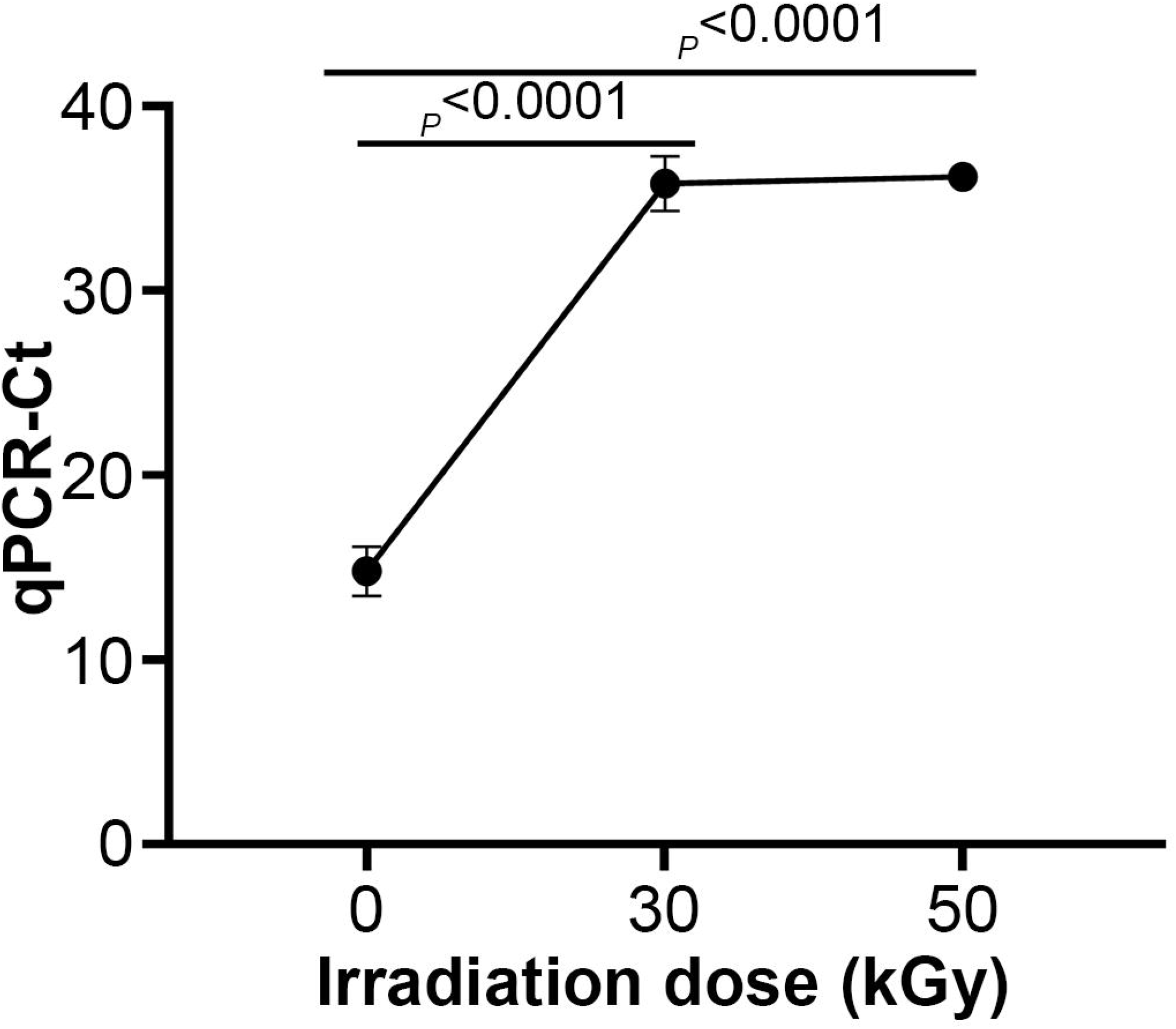
The qPCR Ct values from serially passaged endpoints of both irradiated and non-irradiated samples. The data are x□ ±□ SEM from the 3 replicates. _*P*_*-*values were calculated using a Student t-test.

### Impact of high irradiation doses on the genomic integrity of ASFV

Genome depth of coverage analysis was performed to assess whether high gamma irradiation doses affect the integrity or recoverability of ASFV genomic DNA. All virus samples, including non-irradiated (0 kGy) and irradiated (30 kGy and 50 kGy), showed uniformly high coverage across the top contigs, with overall genome coverage above 99% for all samples (Fig. 5A and Table 3). Average sequencing depth was also highly consistent between samples (69-71X) (Table 3). The coverage-by-position profiles revealed nearly identical patterns across the genome, with shared peaks and troughs observed in all samples (Fig 5A). Irradiation-specific regions of dropout or localized loss of coverage were not observed in both 30 and 50 kGy doses samples. All samples showed high frequency distributions between 60X and 80X sequencing depth, with similar frequency distributions and no shift toward lower depth bins in irradiated samples (Fig 5B).

**Fig. 5.**
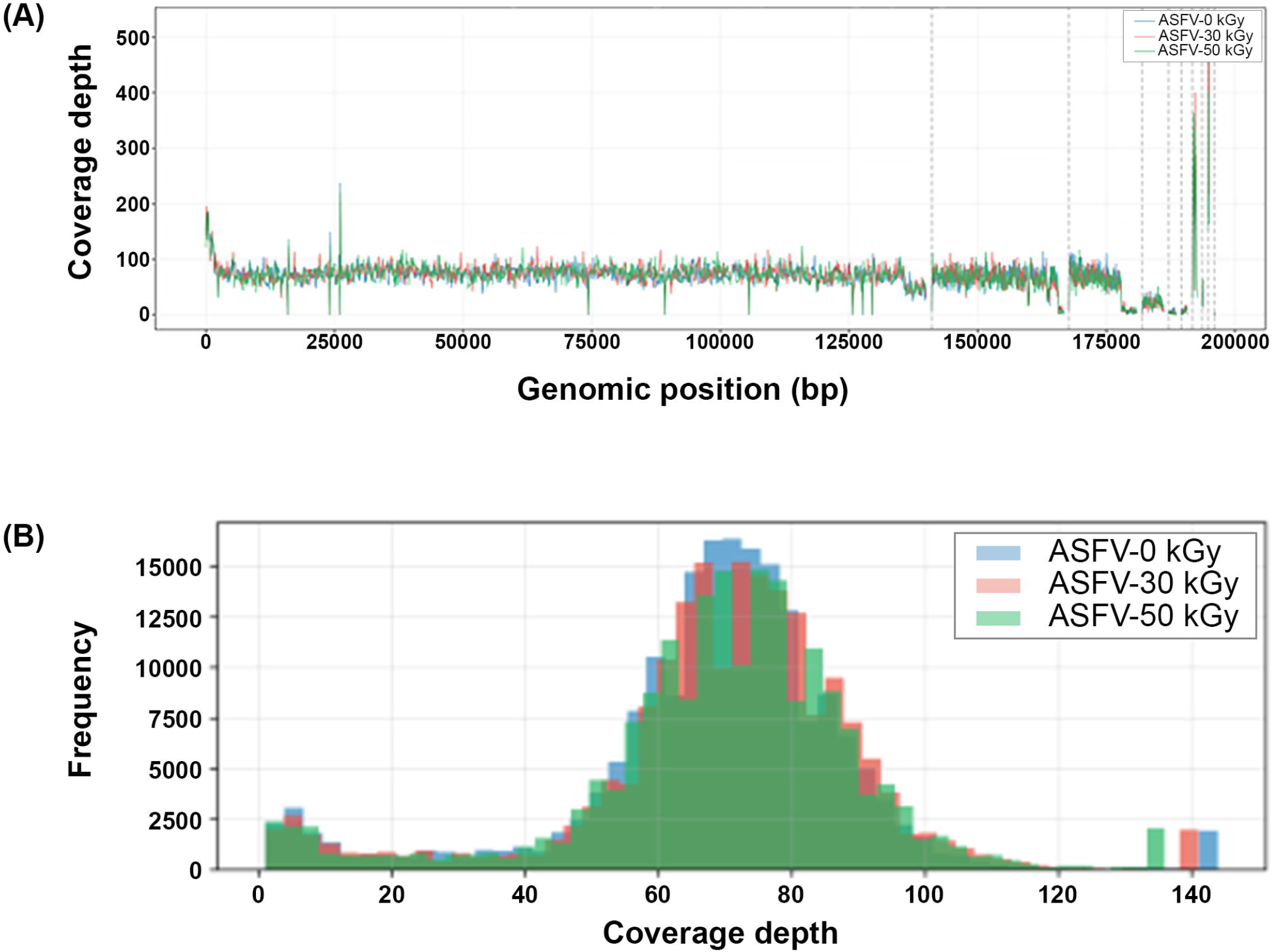
Genome depth-of-coverage analysis between non-irradiated and irradiated samples. **A**. Genome coverage depth map between non-irradiated (0 kGy) and 30 kGy and 50 kGy dose-irradiated samples. **B**. The comparable coverage depth distribution across the genome for non-irradiated (0 kGy) and 30 kGy and 50 kGy dose-irradiated samples.

**Table 3.** Coverage statistics were computed across the genome, including percent coverage and average read depth in covered regions.

| Sample name | Coverage (%) | Avg Depth Coverage (X) |
| --- | --- | --- |
| ASFV-0 kGy | 99.4 | 69.0 |
| ASFV-30 kGy | 99.1 | 70.6 |
| ASFV-50 kGy | 99.0 | 70.1 |

## Discussion

Gamma irradiation is an effective method for inactivation of high-consequence pathogens without disturbing the genome and structural integrity (Elliott et al., 1982; Feldmann et al., 2019). However, complete inactivation of highly pathogenic viruses using gamma irradiation depends on several factors, including virus type, viral genome size, initial titer, sample type, and temperature (Elveborg et al., 2022). Our results demonstrate that gamma irradiation is an effective and reliable method for inactivating ASFV while preserving the molecular integrity required for molecular diagnostic testing. We estimated a ~29 kGy irradiation dose that achieved a SAL of 10□□, the acceptable standard for viral inactivation (Hewitt and Leelawardana, 2014). However, even if the required irradiation dose meets the SAL for virus inactivation, the possibility remains that one live virus is present in the inactivated sample (Hewitt and Leelawardana, 2014). This suggests that the estimated irradiation dose may need to be increased and that further validation for high-consequence viruses, such as ASFV, to ensure complete inactivation. Our *in vitro* viral infectivity validation results provided strong evidence of complete inactivation, as neither the 30 nor 50 kGy irradiated virus samples produced cytopathic effects at any point in the study. This result was supported by Ct values near the lower detection limit in the third blind passage of the irradiated samples, indicating the absence of virus. These results are consistent with previous reports that ASFV is inactivated at a 30 kGy gamma-irradiated dose; however, these studies did not validate the suitability of downstream molecular testing of inactivated samples (McVicar et al., 1982; Boudarkov et al., 2016; Pikalo et al., 2022b).

Higher gamma irradiation doses may induce viral DNA/RNA damage (Leung et al., 2020), which prevents the molecular testing of inactivated samples. A previous study on SARS-CoV-2 showed that qPCR results were not affected upon inactivation using gamma irradiation, but the same study suggested that qPCR sensitivity depends on radiation dose (Leung et al., 2020). Importantly, high-dose irradiation did not compromise molecular detection of ASFV in our study, as supported by qPCR results. Furthermore, sequencing data confirmed that no indels or SNPs (Single Nucleotide Polymorphisms) were present within the qPCR target sequences.

Moreover, ASFV exposed to irradiation doses (30 and 50 kGy) had >99% coverage by WGS suggesting that the integrity of the genomic DNA is maintained following irradiation. This result is supported by the inactivation of viruses through X-ray irradiation, which preserves genome integrity (Afrough et al., 2020). Although gamma irradiation is known to induce nucleic acid damage (Leung et al., 2020), the preserved coverage map suggests that minimal fragmentation occurred following irradiation. This result aligns with our real-time PCR results showing that ASFV lost viral infectivity while maintaining sufficient DNA quality for molecular testing.

Until now, ASFV reference materials and proficiency□testing panels have not been widely available because ASFV is a high□consequence, high□containment pathogen in unaffected countries. In addition, ASFV is a large, complex DNA virus that is difficult to propagate consistently at high titers, further complicating the production of uniform reference stocks. In this study, we generated high□titer ASFV in cell culture and successfully inactivated the virus using gamma irradiation without impact on molecular detection. The resulting inactivated ASFV□cell culture supernatant was produced in compliance with NVSL quality assurance requirements. This inactivated material can be safely distributed to laboratories that do not operate at high biocontainment levels, enabling broader participation in inter□laboratory comparison programs and enhancing national diagnostic harmonization.

Additional validation studies are needed to confirm that a 50 kGy irradiation dose reliably inactivates ASFV across different sample types used for diagnostic testing. Furthermore, we did not validate the immunological properties of the inactivated ASFV sample, as this study utilized only ASFV□infected cell culture supernatant. Additional studies are therefore required to evaluate whether higher irradiation doses preserve structural integrity suitable for serological assays.

Overall, we determined the irradiation dose required to fully inactivate ASFV, demonstrated complete loss of viral infectivity *in vitro*, and confirmed that the resulting inactivated material remains suitable for molecular assays.

## Acknowledgments

We thank Dr. Jennifer Welch for providing the initial stock of ASFV□infected cell culture supernatants. The authors thank Dr. Karthik Shanmuganatham1 and Dr. Randall Levings for his guidance and support, and Patrick Camp for his sequencing expertise. This work would not have been possible without the establishment of the gamma irradiator facility and the federal employees who support it. The findings and conclusions in this publication are those of the authors and should not be construed to represent any official USDA or U.S. Government determination or policy.

